# Specificity and Mechanism of tRNA cleavage by the AriB Toprim nuclease of the PARIS bacterial immune system

**DOI:** 10.1101/2025.02.04.636281

**Authors:** Svetlana Belukhina, Baptiste Saudemont, Florence Depardieu, Tom Lorthios, Tinashe P. Maviza, Alexei Livenskyi, Marina Serebryakova, Maria Aleksandrova, Elizaveta Ukholkina, Nadezhda Burmistrova, Petr V. Sergiev, Marouane Libiad, Sarah Dubrac, Frederic Barras, Yuri Motorin, Virginie Marchand, Gregor Hagelueken, Artem Isaev, David Bikard, Christophe Rouillon

**Author notes:** These authors contributed equally to this study (Svetlana Belukhina, Baptiste Saudemont).

## Abstract

Transfer RNA molecules have been recently recognized as widespread targets of bacterial immune systems. Translation inhibition through tRNA cleavage or modification inhibits phage propagation, thereby protecting the bacterial population. To counteract this, some viruses encode their own tRNA molecules, allowing infection to take place. The AriB effector of the PARIS defence system is a Toprim nuclease previously shown to target the *E. coli* tRNA^Lys(UUU)^, but not a tRNA^Lys(UUU)^ variant encoded by bacteriophage T5. We demonstrate here that the T5-tRNA^Lys(UUU)^ is required but not sufficient to bypass PARIS immunity. Combining tRNA-sequencing, genetics, phage infection and *in vitro* biochemical data, we reveal that the *E. coli* tRNA^Thr(UGU)^, is another prime target of AriB and tRNA^Asn(GUU)^ represents a secondary, yet biologically relevant, target of the PARIS effector. Activated AriB protein cleaves these targets *in vitro*, and the cleavage reaction is not dependent on the presence of specific tRNA modifications. We show that the overexpression of phage T5 tRNA^Lys(UUU)^, tRNA^Thr(UGU)^ and tRNA^Asn(GUU)^ variants is sufficient to inhibit PARIS anti-viral defence. Finally, we propose a model for tRNA recognition by the AriB dimer and provide molecular details of its nuclease activity and specificity.

## Introduction

The central role of tRNAs in protein synthesis makes them prime targets for various toxins inhibiting the growth of bacteria. To compete with neighbours, some bacteria secrete bacteriocins, like colicin E5 and D, which are known to cleave specific tRNAs within their anticodon loop [1]. Multiple Toxin-antitoxin (TA) systems also inhibit translation by targeting tRNAs [2–4]. Although TA systems have been linked to various biological functions, recent studies have highlighted their role as antiphage defence systems, primarily functioning by arresting the growth or killing the host bacteria thereby blocking phage propagation [5]. Targeting tRNAs appears to be a common strategy among antiphage defence systems, as highlighted by the PrrC anti-codon nuclease [6], CRISPR-Cas13 [7], Retron-Eco7 [8], toxSAS-CapRel and FaRel [9,10], RemAIN [11], the PARIS immune system [12] and more [13].

Mechanisms of tRNA inactivation include their modification by nucleotidyltransferases such as MenT [14], by pyrophosphokinases such as CapRel [9], or by acetyltransferases such as TacT [15,16]. Another frequent route of tRNA inactivation involves the cleavage of the anti-codon loop by proteins with a colicin D/E5 nuclease domain, by PIN-domain ribonucleases such as VapC [17,18] and MazF [19], or by HEPN nucleases such as PrrC, RloC [20] and RemN [11]. Other nuclease domains have recently been involved in the degradation of specific tRNAs in the context of anti-phage defence systems. These include the HNH nuclease domain of PtuB in the Retron-Eco7 system [8] or the Toprim domain of AriB in the PARIS system [12]. Toprim domain-containing nucleases play essential roles in DNA replication, repair and recombination [21]. This domain is also found in OLD proteins where it is associated to an ABC ATPase domain [22]. This association has been described across several anti-phage defence systems including the prototypical OLD protein of bacteriophage P2 [23], Gabija [24], retron elements [25], AbiL [26], MADS [27] and the PARIS [28] systems.

PARIS systems are composed of two proteins (AriA and AriB) that are sometimes fused as single polypeptide. AriA is an ABC ATPase which assembles into a hexamer and functions as a sensor, while AriB is a tRNase which contains a Toprim domain and serves as the effector protein. In the absence of phage infection, the AriA hexamer binds to and sequesters three copies of AriB, maintaining them in an inactive state. Upon phage infection, AriA detects specific phage proteins such as Ocr from phage T7 or Ptr1 and Ptr2 proteins from T5 [12,28,29]. These triggers bind to AriA, inducing a conformational change that leads to the release and dimerization of AriB. We recently described how the activated AriB protein cleaves the tRNA^Lys(UUU)^ in the stem of the anti-codon stem-loop between positions 40 and 41, a cleavage site not previously described for other tRNA nucleases [12]. It was proposed that viral tRNAs accumulate mutations to avoid cleavage by the host tRNases [30]. This was confirmed in the case of the T5 tRNA^Lys(UUU)^ which contains two mutations around the AriB cleavage site which are sufficient to provide resistance to AriB cleavage [12].

In this study, we characterize the complete set of AriB Toprim nuclease tRNA targets and describe the mechanism of cleavage in more details. We show that in addition to tRNA^Lys(UUU)^ AriB cleaves tRNA^Thr(UGU)^, and to a lower extent tRNA^Asn(GUU)^ and tRNA^Thr(CGU)^. We demonstrate that cleavage of secondary tRNA targets is essential for PARIS activity, since only overexpression of the full set of phage T5 tRNA variants can rescue PARIS-induced toxicity and inhibit its anti-viral immunity. We further investigate whether base modifications found in tRNAs are required for AriB target recognition, and the RNA sequence determinants of this recognition. Finally, we employ Alphafold3 (AF3) to propose a model of how two catalytic sites within the AriB dimer engage the asymmetrical tRNA substrate leading to a unique cleavage site. Altogether our results shed light on the molecular mechanism underlying tRNA cleavage by the Toprim domain of AriB, a protein domain with homologues found across many uncharacterized anti-phage defence systems.

## Results

### Landscape of tRNA cleavage by AriB

The tRNA^Lys(UUU)^ was previously identified as a target of PARIS based on the observation that the T5 tRNA^Lys(UUU)^ could rescue the infection of a deletion variant of T5 (T5_123_) in which 11 of the 24 tRNAs were lost (deletion spanning positions 29,191-32,442, according to GenBank assembly AY543070.1). Since T5_123_ still carries 13 tRNA genes, we could not exclude that AriB also cleaves additional *E. coli* tRNAs, and that other phage tRNAs contribute to countering PARIS immunity. To test this hypothesis, we expressed the T5-tRNA^Lys(UUU)^ in the presence of PARIS during infection by T5_Mos_ another T5 deletion variant (30,661-38625nt) that lost a slightly different set of 17 tRNA genes (29,191-30,661nt) [12]. Although T5 tRNA^Lys(UUU)^ expression completely rescued T5_123_ infectivity, this was not sufficient in the case of T5_Mos_ (figure 1A). Thus, tRNA^Lys(UUU)^ from T5 is necessary, but not sufficient to counteract the effect of AriB activation, suggesting the existence of additional tRNA targets.

**Figure 1.**
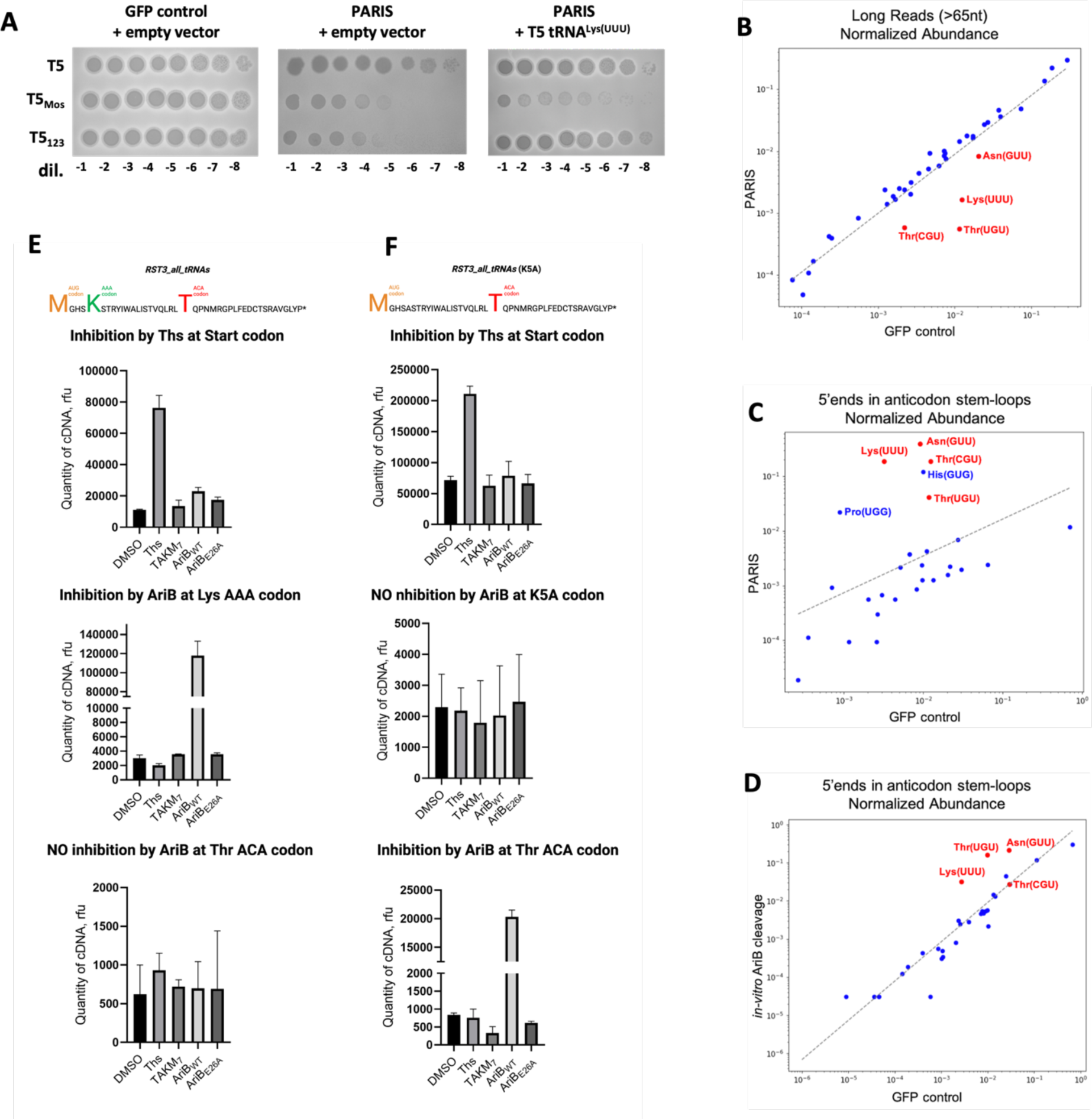
Landscape of AriB-mediated tRNA cleavages. **A.** Plaque assays were conducted using cells without (GFP control) or with PARIS in the presence of empty vector (pBAD) or pFD287 (paraBAD T5 tRNA^Lys(UUU)^) against T5WT, T5Mos and T5123. Overexpression of T5 tRNA^Lys(UUU)^ in PARIS cells from pFD287 plasmid, with the tRNA^Lys(UUU)^ of phage T5 under the control of the araBAD promoter inducible by L-ara (0.2%), rescues infection by phage T5123 but not by T5Mos. **B-D.** Scatter plots of normalized tRNA abundances. Axes represent the normalized abundance of tRNAs in the respective samples. Blue dots indicate the majority of tRNAs, while red dots highlight specific tRNAs of interest: Lys(UUU), Thr(UGU), Thr(CGU), and Asn(GUU). The dashed gray line shows the linear regression fit in log-log space. **B.** tRNAs with mapped reads ≥65 nt from GFP control and PARIS (*in vivo* cultures), sequenced using a custom protocol (see methods). **C.** Small tRNAs (<45 nt) with 5ʹ ends mapped to the anticodon stem-loop, from GFP control and PARIS (*in vitro* cultures), sequenced using the NEBNext kit. **D.** Small tRNAs (<45 nt) with 5ʹ ends mapped to the anticodon stem-loop, from untreated GFP control and *in vitro* AriB cleavage samples, sequenced using the NEBNext kit. **E-F** Frequency of ribosome pausing on specific codons, estimated by toeprinting analysis with substrates *RST3_all_tRNAs* (**E**) and *RST3_all_tRNAs (K5A)* (**F**) mRNA templates translated in the presence of AriB WT or E26A Toprim mutant. Sequences of mRNA templates are provided at the top. **‘**Ths’ represents a control reaction with thiostrepton, which stalls ribosomes at the start codon. Signals from the start (AUG), lysine (AAA), and threonine (ACA) codons were calculated as the mean area under the curve from the capillary electropherograms carried in triplicates, representative electropherograms are presented in the (figure S1D-E).

To determine the comprehensive set of AriB targets, we aimed to sequence tRNAs extracted 30 min following the induction of T7 Ocr in *E. coli* cells expressing PARIS or a GFP control (figure 1B-D; figure S1A). We first implemented a custom protocol that yielded a high number of reads covering more than 80% of the tRNA length. These reads served as a proxy to calculate the abundance of full-length, functional tRNAs. This analysis revealed a depletion of tRNA^Lys(UUU)^, tRNA^Thr(UGU)^, tRNA^Asn(GUU)^, and tRNA^Thr(CGU)^ following the induction of PARIS and its trigger (figure 1B). We were however unable to clearly identify tRNA cleavage products in this dataset. tRNAs contain numerous modified bases that can cause the reverse transcriptase enzyme to stall, resulting in shorter cDNAs compared to their templates [31,32]. Our custom protocol was adding sequencing adapters to cDNAs, resulting in a high coverage sequencing of these short cDNAs which can obfuscate cleavage products. To address this, we used a commercial kit that ligates sequencing adapters to RNA ends prior to reverse transcription, ensuring PCR amplification is limited to fully reverse-transcribed products. Using this data, analysis of 5’ ends within the anti-codon stem-loop identified probable cleavage products for tRNA^Lys(UUU)^, tRNA^Thr(UGU)^, tRNA^Asn(GUU)^, tRNA^Thr(CGU)^, tRNA^His(GUG)^, and tRNA^Pro(UGG)^ (figure 1C). Some of the tRNAs detected with this method could be minor targets of AriB which are not cleaved efficiently enough to show a depletion signal from the pool of full length tRNAs. Notably, a cleavage between position 40 and 41 could be detected for most of these tRNAs (figure S1B) as previously described for the tRNA^Lys(UUU)^. Some of these cleavage products could also be indirect consequences of translation arrest after AriB activation. To further validate the targets of AriB we performed an *in vitro* cleavage assay in which *E. coli* tRNAs were incubated with activated AriB protein (figure S1A). Sequencing enabled the detection of cleavage events in the tRNA^Lys(UUU)^, tRNA^Thr(UGU)^ and tRNA^Asn(GUU)^ (figure 1B). Altogether, the main targets of AriB appear to be tRNA^Lys(UUU)^, tRNA^Thr(UGU)^ and tRNA^Asn(GUU)^ with some weaker activity on tRNA^Thr(CGU)^.

To test whether tRNA cleavage by AriB would lead to ribosomal stalling at the expected codons, we performed toeprinting analysis. In this assay, mRNAs undergoing *in vitro* translation are subjected to reverse transcription with fluorescein-labelled primer. The reverse-transcriptase is blocked by stalled ribosomes, mapping their position (figure S1C). To unambiguously identify codons affected by AriB tRNA cleavage, we designed an mRNA template that contained a codon for each known *E. coli* tRNA species [31]. Toeprinting revealed that presence of the AriB WT, but not AriB E26A Toprim mutant, resulted in the accumulation of ribosomes stalled at the lysine AAA codon (figure 1E, figure S1D). To identify secondary AriB targets, we mutated the lysine codon in the mRNA to an alanine codon and repeated toeprinting, which revealed ribosome pausing at the downstream threonine ACA codon, recognized by tRNA^Thr(UGU)^, although milder compared to the lysine codon (figure 1F, figure S1E). Mutating both AAA and ACA tRNA codons to alanine codons did not reveal significant ribosome pausing at other sites. These *in vitro* results confirm that AriB cleavage of tRNA^Lys(UUU)^ and tRNA^Thr(UGU)^ blocks translation at the expected codons. Cleavage of the tRNA^Asn(GUU)^ was likely not efficient enough to substantially stall ribosomes in this assay.

### Multiple T5 tRNAs are needed to rescue AriB toxicity and inhibit PARIS immunity

We decided to take the validated targets tRNA^Lys(UUU)^ and tRNA^Thr(UGU)^, as well as the tRNA^Asn(GUU)^ detected by sequencing to test whether co-expression of T5 homologs of those tRNAs is sufficient to inhibit AriB toxicity. To do so we used cells co-expressing PARIS and Ocr and induced T5 tRNAs expression at different arabinose concentrations. When Ocr is induced, cells expressing PARIS die as a result of AriB activation. Expression of tRNA^Lys(UUU)^ alone did not rescue PARIS toxicity, while co-expression with tRNA^Thr(UGU)^ increased cell survival. Growth could however be almost completely rescued by the co-expression of the three T5 tRNA^Lys(UUU)^/tRNA^Thr(UGU)^/tRNA^Asn(GUU)^ (figure 2A). In conditions of enhanced expression (2% arabinose induction), co-expression of the two tRNA^Lys(UUU)^/tRNA^Thr(UGU)^ was sufficient to restore cell growth (figure 2A).

**Figure 2.**
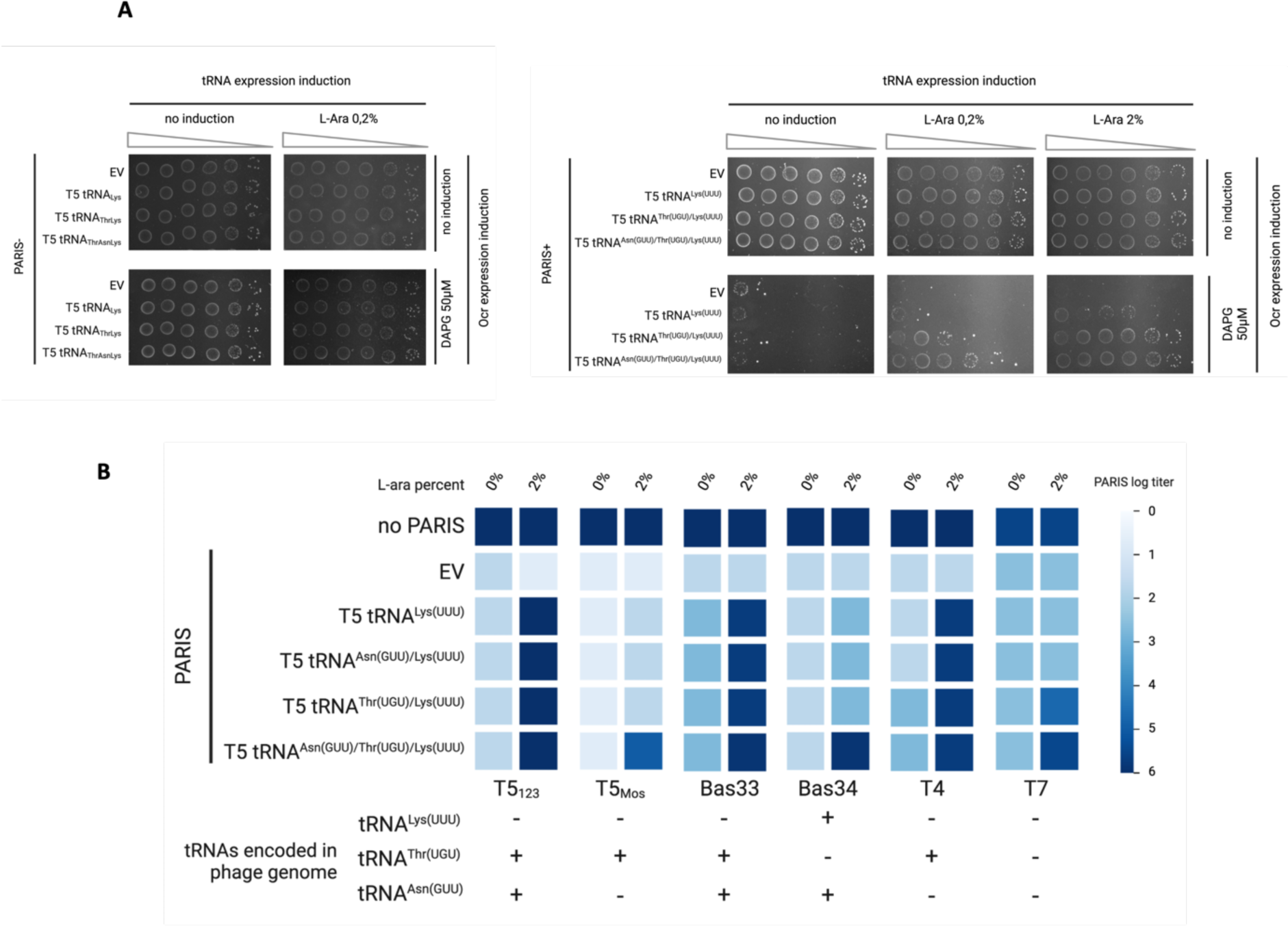
Expression of viral tRNAs reduces AriB toxicity and restores phage infectivity. **A.** Phage T5 tRNA^Lys(UUU)^/tRNA^Thr(UGU)^/tRNA^Asn(GUU)^ expression can rescue PARIS toxicity triggered by the expression of the phage T7 Ocr from an arabinose (Ara) inducible pBAD promoter. **B.** Heatmap of PARIS defence against a panel of phages. Color intensity indicates plaque forming units (PFUs) on a log10 scale. A panel below shows whether phages encode their own variant of the tested tRNAs. Representative plates used to build the heatmap are shown in (figure S2).

We then tested whether the expression of one or several of phage T5 tRNAs would rescue the infection of phages susceptible to the PARIS defence system. We performed an EOP assay with the same phages in cells expressing T5 tRNA^Lys(UUU)^, T5 tRNA^Lys(UUU)^/tRNA^Thr(UGU)^, T5 tRNA^Lys(UUU)^/tRNA^Asn(GUU)^,or T5 tRNA^Lys(UUU)^/tRNA^Thr(UGU)^/tRNA^Asn(GUU)^ (figure 2B, figure S2). As previously observed, infectivity of T5_123_ was rescued by expressing only T5 tRNA^Lys(UUU)^, while infectivity of T5_Mos_ was rescued only in conditions of three tRNAs overexpression even though it encodes its own copy of tRNA^Thr(UGU)^. We further noticed that T5 relatives, phages Bas33 and Bas34, which are naturally sensitive to PARIS defence, encode either tRNA^Thr(UGU)^ or tRNA^Lys(UUU)^ but not both (figure 2B). Expression of tRNA^Lys(UUU)^ was sufficient to rescue Bas33 infection, while rescue of Bas34 required all three tRNAs. We next showed that phage T4, which encodes tRNA^Thr(UGU)^, can be rescued by expression of T5 tRNA^Lys(UUU)^. Finally, infection of the T7 phage, which carries no tRNAs, could be rescued by the co-expression of T5 tRNA^Lys(UUU)^/tRNA^Thr(UGU)^ or the 3 T5 tRNAs (tRNA^Lys(UUU)^/tRNA^Thr(UGU)^/tRNA^Asn(GUU)^).

Together, these results demonstrate that various viral tRNA contribute to the inhibition of PARIS immunity and confirm previous observations of AriB tRNA specificity but raise additional questions regarding the mechanism of escape. It is somewhat surprising that the T5 tRNA^Asn(GUU)^ was required to rescue infection of T5_Mos_, but not T4 and T7, lacking this tRNA. At the same time, expression of tRNA^Thr(UGU)^ from plasmid was required to rescue T5_Mos_ infection, and expression of T5 tRNA^Asn(GUU)^ was required to rescue Bas34 infection, despite those tRNAs being encoded in the respective phage genomes. These differences could highlight the varied sensitivity of phages to the depletion of certain tRNAs. It is also possible that in addition to rescuing translation, viral tRNAs act as inhibitors of AriB when overexpressed, maybe through competition with the *E. coli* tRNAs.

### tRNA modifications are not essential for specific AriB cleavage

We then investigated the importance of tRNA modifications for AriB recognition and cleavage. In *E. coli*, there are 43 known tRNA modifications, some of which are required for cleavage by anti-codon nucleases. PrrC, which cleaves the tRNA^Lys(UUU)^ of *E. coli* between positions U_35_ and U_36_ requires cyclic threonylcarbamoyl adenosine (ct6A) at position 37 [33]. The bacterial toxin colicine E5 targets tRNAs with queuosine modification in the wobble position of their anticodon and the colicine D preferentially targets modified tRNA^Arg(CGN)^ [1]. In *E. coli* tRNA^Lys(UUU)^, 10 nucleotides are modified with 7 different types of modifications [34]. Using Modomics [35], we revealed that *E. coli* tRNAs Lys(UUU), Thr(UGU), Asn(GUU) and Thr(CGU) contain 13 modifications distributed on similar nucleotides (figure S3A).

To assess the importance of these modifications for PARIS activity, we used *E. coli* strains that cannot produce certain modifications, including pseudouridines Y39 (mutant *ΔtruA*), Y55 (mutant *ΔtruB*), ct6A37 (*ΔtcdA*)[36] and mnm^5^s^2^U34 in the wobble position (*ΔmnmA* mutant)[37]. For all mutants tested, we found that PARIS activation triggers cell death at the same level as the wild-type strain (figure 3A).

**Figure 3.**
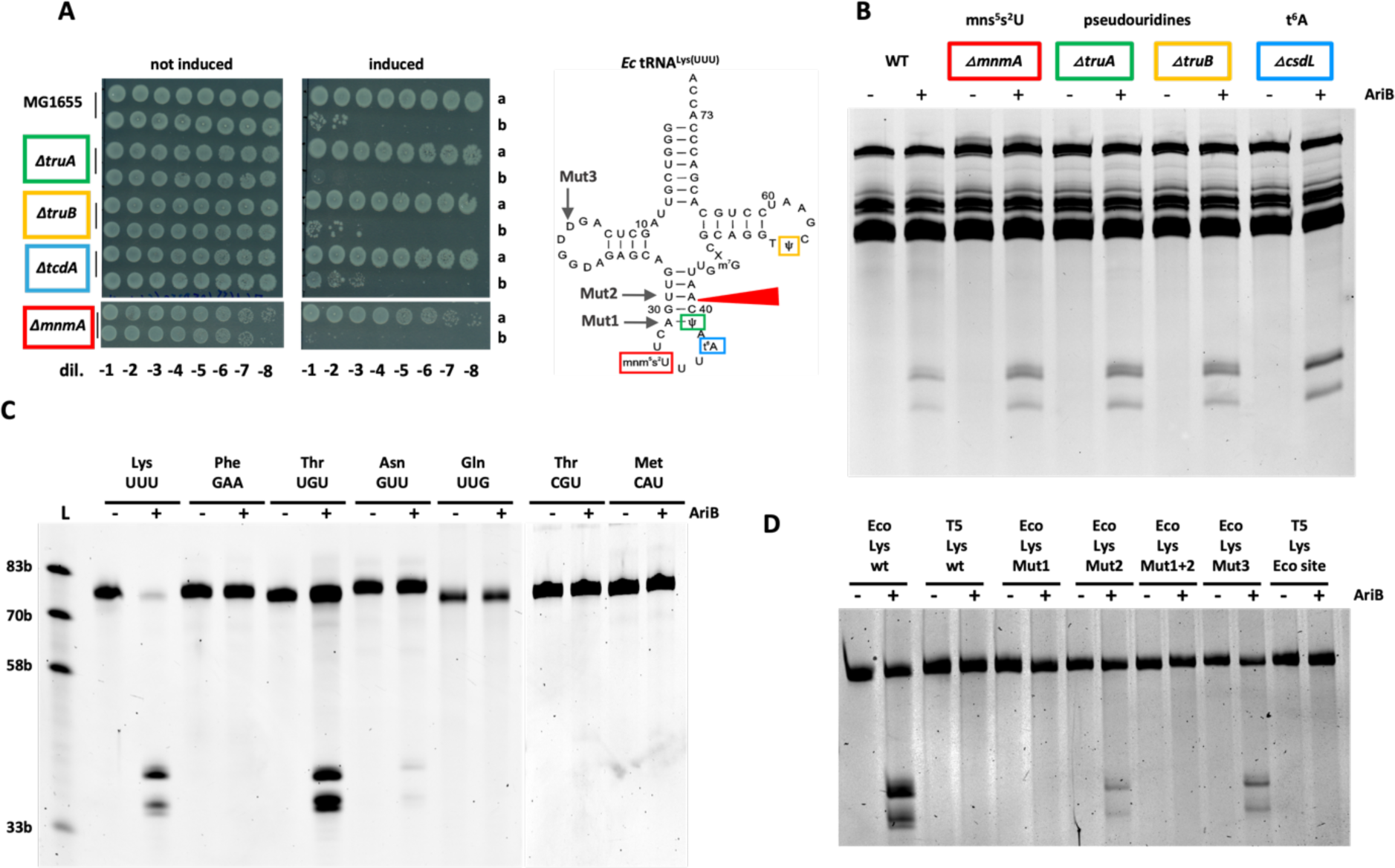
AriB does not require tRNA base modifications for cleavage. **A.** AriB toxicity was triggered by inducing the expression of T7 Ocr under the control of a PPhlF promoter inducible by DAPG (50 µM) in *E. coli* with different deletions of genes implicated in base modifications Bacteria carried either the PARIS system (b) or a GFP control vector (a). **B.** *in vitro* AriB cleavage assay on tRNAs extracted from the four deletion strains or the WT strain. **C.** Activity of AriB on specific *in vitro* transcripts corresponding to *E. coli* tRNAs. **D.** Activity of AriB on *E. coli* and T5 tRNA^Lys(UUU)^ and mutants; point mutations are shown in (figure S3D).

We further confirmed that activated AriB can cleave tRNAs extracted from four *E. coli* mutant strains (*ΔtruA*, *ΔtruB*, *ΔtcdA* and *ΔmnmA*) *in vitro* (figure 3B). Those data suggest that AriB, in contrast to PrrC or tRNA-targeting colicines, does not require base modifications to cleave its target. To further validate this, we tested the activity of purified AriB on an *in vitro* transcribed tRNA^Lys(UUU)^, lacking all modifications. In accordance with recently published results [29], we were able to see a strong cleavage of the transcript at the expected nucleotide position (figure S3B). In contrast, AriB shows no cleavage of the control *E. coli* tRNA^Phe(AAA)^ (figure S3B). Thus, base modifications are not required for AriB cleavage *in vitro*, but it remains possible that base modifications contribute to the interaction of AriB with tRNA targets *in vivo*.

### Sequence specificity determinants of AriB tRNase

To gain further insights on the tRNA specificity, we challenged AriB with a larger set of *in vitro* transcribed tRNAs. The tRNA^Thr(UGU)^ substrate was efficiently cleaved by AriB, although not as efficiently as tRNA^Lys(UUU)^ (figure 3C). The tRNA^Asn(GUU)^ demonstrated significantly reduced cleavage, whereas tRNA^Thr(CGU)^ cleavage was barely visible on gel and cleavage of other control tRNA (i.e., tRNA^Gln(UUG)^ and tRNA^Met(CAU)^) was not detected (figure 3C). Interestingly, the sequence UAAU present near the tRNA^Lys(UUU)^ cleavage site U_36_AAUC|A_41_ (cleavage marked with |), is conserved in the other major target tRNA^Thr(UGU)^ and in both minor targets tRNA^Asn(GUU)^ and tRNA^Thr(CGU)^ (figure S3C Top). Regarding the cleavage site, whereas tRNA^Thr(UGU)^ has the same as tRNA^Lys(UUU)^, the tRNA^Asn(GUU)^ has a substitution at position 41 U_36_AAUC|C_41_ and tRNA^Thr(CGU)^ has substitutions on both nucleotides 40-41 at the cleavage site U_36_AAUG|C_41_ (figure S3C Top). We previously noticed that T5 tRNA^Lys(UUU)^, which is resistant to cleavage by AriB [12], had two substitutions we called Mut1 (U_39_>A) and Mut2 (A_41_>C). Here, using *in vitro* transcribed tRNA we confirm that the T5 tRNA^Lys(UUU)^ is not cleaved by AriB (figure 3D). The mutation Mut1 (U_39_>A) introduced to the *E. coli* tRNA^Lys(UUU)^ transcript completely abolishes AriB cleavage, whereas Mut2 (A_41_>C) substantially inhibits it (figure 3D, figure S3D). To investigate whether the recognition of the deduced AriB site is sufficient to promote tRNA cleavage, we reintroduced the *E. coli* cleavage site into the T5 tRNA^Lys(UUU)^ (U_36_AAACC_41_ > U_36_AAUCA_41_), but this new hybrid tRNA was not cleaved by AriB (figure 3D). This suggest that in addition to recognition of the cleavage site, AriB forms additional base specific interactions with tRNAs at positions distal to the anticodon loop, which are different between T5 and *E. coli* tRNA^Lys(UUU)^.

### The two catalytic sites of the AriB dimer interact with distinct tRNA elements

To better understand how AriB distinguishes between target and non-target tRNAs and why the AriB dimer produces a single nick at the cleavage site, we predicted the structure of the AriB dimer in complex with the *E. coli* tRNA^Lys(UUU)^ using AF3 [38]. This resulted in a structural prediction with a high degree of overall confidence (pTM 0.92, ipTM 0.89). Especially the dimerization interface of the AriB protein is predicted with high confidence values, while local features of the tRNA are not predicted as confidently according to the predicted local difference distance test (pLDDT) (figure S4A). Nonetheless, the overall structure and placement of the tRNA with respect to the protein dimer was identical between many runs with very small errors in the predicted Alignment Error (pAE) matrices (figure S4A right). We were intrigued by this prediction of a symmetric protein dimer interfacing with an asymmetric RNA molecule (figure 4A). Strikingly, in the prediction, one of the AriB monomer nuclease catalytic site is clearly positioned in front of the nucleotides constituting the cleavage site (C_40_-A_41_). This increased our confidence in the AF3 model, and we decided to analyse its features in more detail and to design experiments to validate the model. Interestingly, two lysine (K60 and K64) located on a small alpha helix seem to hold the anticodon loop above the cleavage site, like tweezers (figure 4B left, PSE file). Indeed, mutation of these lysine to alanine residues abolished defence against phage T7 by PARIS (figure 4B right). This experimental observation agrees with the AF3 structure suggesting a stabilisation of the anticodon loop by the lysines K60 and K64 of the AriB monomer that displays cleavage. We then looked at the second catalytic site, wondering why AriB dimer introduces only a single nick. The second catalytic site faces two nucleotides (U_16_-U_17_) of the D-Loop, that are represented by Dihydrouridines in the native tRNA. In contrast to C_40_-A_41_ in the active catalytic site, AF3 models the U_16_-U_17_ nucleotides in a flipped position and interacting with positively charged residues of AriB (R28-U_16_ and R236-U_17_), in an arrangement that is apparently not compatible with a cleavage reaction (PSE file). Of note, the nucleotide U_16_ is present in *E. coli* tRNA^Lys(UUU)^ and tRNA^Thr(UGU)^ but is substituted by a cytosine in T5 (figure S3D). Introducing the Mut3 (U_16_>C) substitution in *E. coli* tRNA^Lys(UUU)^ significantly impaired AriB cleavage (figure 3D). This suggests that the non-active AriB monomer makes base specific interactions with the D-loop.

**Figure 4.**
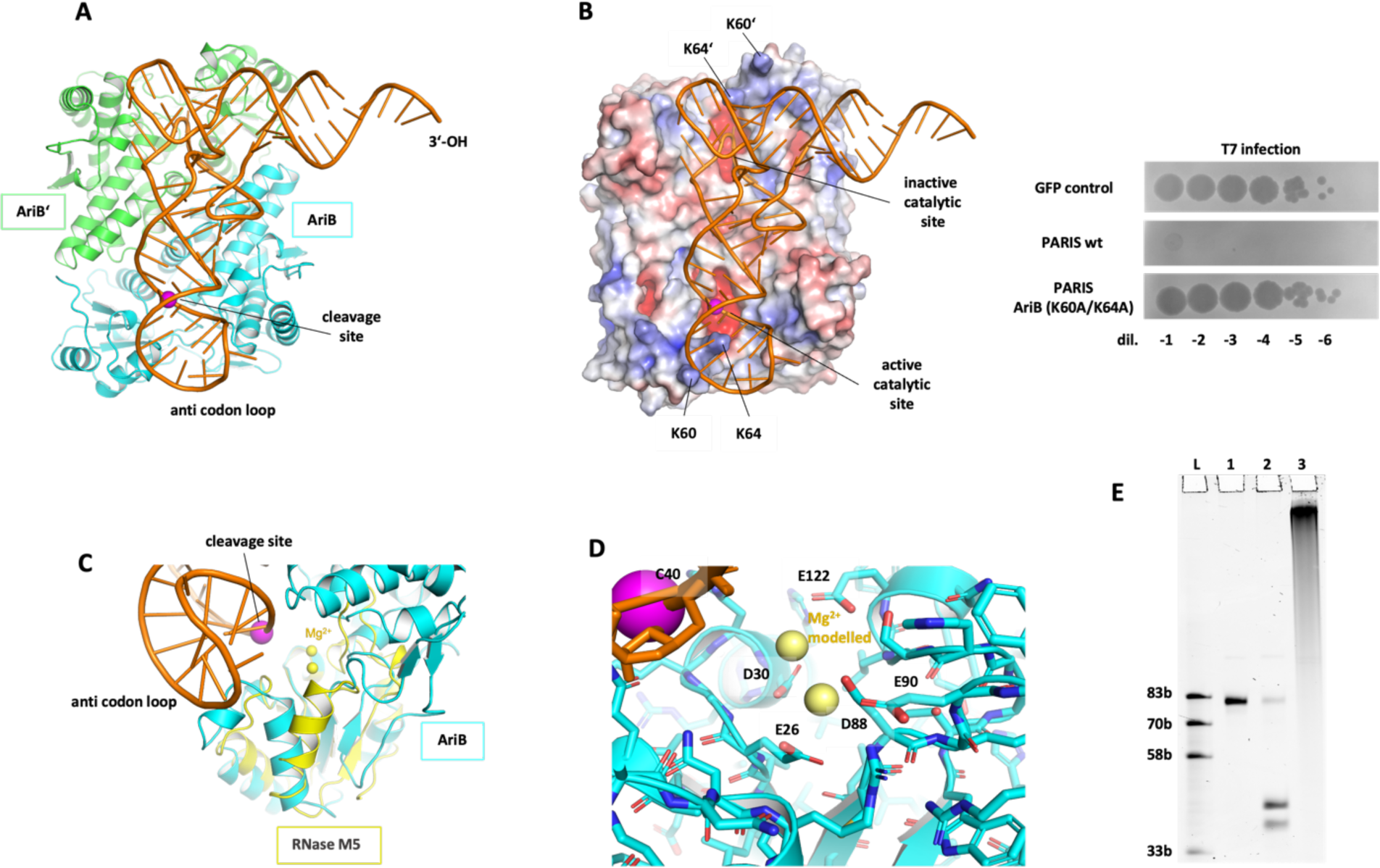
Structure and mechanism of AriB. **A.** Overall structure of AriB dimer bound to *E. coli* tRNA^Lys(UUU)^ generated with AF3 and showing the monomers AriB and AriB’ in light blue and light green. The cleavage site is marked by a magenta sphere **B.** Left. View of the electrostatic surface of the AriB dimer bound to the *E. coli* tRNA^Lys(UUU)^ showing the two lysines K60/K64 stabilizing the anti-codon loop on the AriB monomer with the active catalytic site. Right. Mutations predicted to stabilize the anticodon loop (K60A K64A) prevent PARIS-mediated defence. Efficiency of plaquing of phage T7 on *E. coli* MG1655 carrying the WT or mutated PARIS system. **C.** Superposition of the M5 active site with the AriB active site in complex with the *E. coli* tRNA^Lys(UUU)^. **D.** Active catalytic site of AriB represented with two Mg2+ ions modelled **E.** Incubation of AriB cleavage products with Poly(A)polymerase ran on a 12% denaturing gel, 1: ssRNA ladder, 2: 100nM transcript tRNA^Lys(UUU)^ 3: same as 1 + 25nM AriB dimmer + 1mM Mg^2+^, 4: same as 3 with Poly(A)polymerase treatment post cleavage.

### Cleavage mechanism of AriB

The catalytic site of AriB, as we proposed earlier based on the cryoEM structure of the PARIS complex [12], should be a two metal ions ribonuclease. We looked for structural homologs of AriB and found that the AriB catalytic site has structural similarity to the M5 Toprim nuclease, which has a bi-metal cluster with a shared hydroxyl in its active site (figure 4C) [39]. The key amino acid side chains to coordinate the metal cluster are conserved between M5 RNase and AriB, supporting the notion that AriB is a two-metal ion ribonuclease (figure 4D).

We tested the metal dependency of AriB, and found Mg^2+^, Mn^2+^, and Co^2+^ supporting AriB-mediated tRNase activity (figure S4B). Interestingly, the Toprim OLD nuclease has also been reported to be a two-metal ion enzyme activated by Co^2+^ on top of Mg^2+^, Mn^2+^, and Ca^2+^ for DNA cleavage [23]. In the cleavage reaction of a two-metal ions nuclease, one metal ion activates water into a nucleophile to attack the nucleic acid phosphate group, breaking the phosphodiester bond, while the other stabilizes the transition state. This reaction is expected to leave a 5’ phosphate and a 3’ hydroxyl tRNA ends after cleavage as previously depicted [40]. We confirmed the presence of a 3’-OH end by incubating the *in vitro* cleaved tRNA^Lys(UUU)^ transcript in presence of ATP and Poly(A) polymerase, which showed the strong shift expected after polyadenylation of a 3’-OH RNA substrate (figure 4E). We further identified cleavage products using mass-spectrometry. AriB reaction products were treated with T1 RNase, which should produce a 14-bp fragment containing the AriB cleavage site in case of the intact tRNA^Lys(UUU)^, while AriB-mediated nicking between 40th and 41st nucleotides should result in the production of two smaller fragments after T1 cleavage (figure S4C). Indeed, we confirmed the production of the smaller fragments after treatment with AriB WT but not with the AriB E26A mutant. Identified masses of these fragments support the formation of 3’-OH, and 5’-PO_4_ products (figure S4D), in line with AriB being a two-metal ions endoribonuclease.

## Discussion

Toprim nuclease domains have now been found in diverse anti-phage defence systems were their function remains largely unknown. Here, we investigated the mechanism of the AriB Toprim nuclease from the PARIS bacterial immune system. Upon detection of phage trigger proteins by the AriA ATPase sensor, AriB is released as a dimer to cleave specific host tRNAs, disrupting translation and inhibiting phage replication. Using a combination of tRNA sequencing, *in vitro* cleavage, and ribosome toeprinting assays, we revealed the full set of tRNAs targeted by the AriB nuclease of *E. coli* strain B185. AriB activation leads to the depletion of the previously described tRNA^Lys(UUU)^, but also of tRNA^Thr(UGU)^, tRNA^Asn(GUU)^ and tRNA^Thr(CGU)^. All these tRNAs were found to be cleaved between position 40 and 41, in the stem of the anti-codon stem-loop, with tRNA^Lys(UUU)^ and tRNA^Thr(UGU)^ being the most efficiently targeted tRNAs. We confirmed the biological significance of these tRNA targets by showing that the joint expression of three tRNA variants (tRNA^Lys(UUU)^, tRNA^Thr(UGU)^, tRNA^Asn(GUU)^) from bacteriophage T5 are necessary and sufficient to rescue *E. coli* growth upon PARIS activation. These phage tRNAs all carry nucleotide substitutions at or around the cleavage site of AriB (figure S3C bottom), and we show for tRNA^Lys(UUU)^ that mutations at this location indeed abrogate AriB cleavage. Interestingly, the infection of phages blocked by PARIS could sometimes be rescued by the overexpression of just one or two of these tRNAs. This could in some cases be explained by the presence of a variant of the missing tRNA in the genome of the rescued phages, but not always. These experiments reveal differences in the susceptibility of phages to tRNA depletion by PARIS, which remain to be investigated.

Our results, together with the recent discovery of other anti-phage systems targeting tRNAs, support the hypothesis that phages are under selective pressure to carry diverse tRNAs to overcome bacterial immunity[12,30]. These viral tRNAs may rescue the depletion of the host tRNAs or could also compete with host tRNAs and form non-productive complexes with immune effectors, ensuring that tRNAs are still available for the translation of phage proteins. Other phages may also employ different strategies to counteract tRNA cleavage. For example, the T4 phage employs a kinase (Pnk) and ligase (Rnl1) to repair tRNAs cleaved in the anticodon loop by the PrrC defence system [41].

Using AF3 predictions, we were able to propose a model according to which AriB functions as a symmetric dimer that binds to asymmetric tRNA substrates, resulting in single-site cleavage. This substrate asymmetry forces the two monomers of AriB to adopt distinct interactions with the tRNA. One monomer’s catalytic site precisely nicks the anticodon stem, while the second catalytic site can apparently not form a productive substrate complex with the tRNA. Clearly, experimental structures are needed to validate and fully understand this intriguing interaction. Experiments with phage-host chimeric tRNA transcripts suggest that in addition to base specific contacts in the anticodon stem and the D-loop, AriB likely also makes specific contacts in other parts of the tRNA, as changing the T5 tRNA^Lys(UUU)^ sequence for that of the *E. coli* tRNA^Lys(UUU)^ in anti-codon stem loop did not restore cleavage of this tRNA.

A close structural homolog of AriB’s catalytic site is the M5 Toprim nuclease, which is involved in ribosomal RNA maturation and cleaves a double-stranded RNA (dsRNA) helix [42]. Unlike AriB, M5 acts as a monomer, cleaving both RNA strands sequentially by repositioning its nuclease domain for the second cleavage [39]. These differences highlight how Toprim domains have evolved different properties, but interestingly the first cleavage site of M5 is identical to that of AriB (5’-CA-3’). Consistently with M5, we show that AriB is a two-metal ion Toprim nuclease leaving 5’ phosphate and 3’ hydroxyl ends after cleavage. Its activity depends on the presence of metal ions such as Mg²⁺ or Mn²⁺, which are crucial for catalysis. Interestingly, this metal dependency is not absolute, as Co²⁺ can substitute for Mg²⁺. The ability to use Co²⁺ was also noted for other nucleases, including Toprim nucleases [23].

The catalytic activity of other Toprim domain proteins involved in anti-phage defence has also been investigated. This includes the OLD nuclease from bacteriophage P2, which has been studied since the 70’s for its role in blocking infection by phage lambda [43]. Biochemical characterization of the OLD nuclease revealed exonuclease activity on double-stranded DNA as well as nuclease activity on single-stranded DNA and RNA [44]. Similar activities, including nicking, were more recently found for other variants of this protein and their crystal structure determined [23,45–47]. In addition to OLD, the Gabija bacterial immune system, also known as class 2 OLD, also carries a Toprim domain in the GajA protein (figure S4E). The Gabija complex was described to cleave double stranded DNA *in vitro* by action of a GajA dimer nicking each DNA strand at a specific sequence motif [24,48,49]. Further work will be needed to understand the molecular basis of the differences in substrate specificity between AriB, OLD and GajA. We also cannot exclude that some of these proteins might behave differently *in vivo* and *in vitro* where parameters such as salt and metal concentrations could produce artifacts. An old manuscript on the P2 OLD system suggested that this protein actually targets *E. coli* tRNAs [50]. While this question has not been reinvestigated since the 70’s, confirming these results with modern techniques would help reconcile our findings on AriB with the mechanism of other OLD nucleases.

Finally, several aspects of the AriB mode of action are still not understood. This includes how AriB discriminates between target and non-target tRNAs. It will be interesting to understand whether the list of targets is conserved across different PARIS systems (Burman et al 2034), and possibly across other Toprim nucleases, or whether AriB is under selective pressure to diversify its substrate specificity to escape phage countermeasures. Another aspect relates to the activation mechanism of AriB. Our previous data supports the hypothesis that AriB is activated by AriA, and not simply released, after a phage protein is sensed [12]. The mechanism of this activation remains to be elucidated. Finally, could tRNA-derived small RNA (tsRNA)[51] generated by AriB be involved in other downstream reactions? We still do not understand the mechanistic consequences of tRNA cleavage by AriB. While the depletion of the pool of specific tRNAs will naturally lead to translational arrest, we previously showed that bacteria lose their membrane integrity within 20min of AriB activation [12]. The chain of events leading to this, and whether bacteria might recover or not from AriB activation also remains to be investigated.

## Supporting information

PSE file Structure

## Contributions

Conceptualization, C.R., A.I. and D.B.; methodology, S.B., B.S., C.R., F.D., T.P.M., A.L. and D.B.; Molecular cloning, microbiology and phage methods, S.B., M.A., F.D. and E.U.; tRNAs preparation, sequencing and analysis, B.S., V.M., Y.M. and D.B.; Toe print analysis, T.P.M., and N.B.; Mutants for tRNA modification study, F.D., S.D, M.L. and F.B.; *In vitro* AriB activity, C.R., B.S., T.L.; Structural study, T.L., G.H. and C.R.; Mass-spectrometry, A.L. and M.S.; resources, A.I., D.B. and P.S.; writing—original draft preparation, C.R., D.B. and A.I.; writing—review and editing, C.R., A.I., B.S. and D.B.; visualization, S.B., T.P.M., and A.L. All authors have read, commented and agreed to the published version of the manuscript.

## Acknowledgments

We thank Valérie de Crécy-Lagard for discussion on tRNA modifications and Blake Wiedenheft for giving feedback on the manuscript.

## Data availability statements

tRNA sequencing data: BioProject ID PRJNA1213714

## Materials and Methods

### tRNA extraction and sequencing and analysis

The preparation of enriched small RNA fractions from total RNA samples has been done as previously described [12].

Small RNA libraries focusing on long reads were prepared using a custom protocol. Up to 200 ng of small RNAs were dephosphorylated using Quick CIP (NEB) in 1X rCutSmart buffer at 37°C for 15 min, followed by purification with the Monarch® Spin RNA Cleanup Kit (NEB). For 3’ adapter ligation, 180 ng of dephosphorylated RNA was ligated to a pre-adenylated adapter (SB104) using T4 RNA Ligase 2, truncated KQ (NEB), in 1X T4 RNA Ligase Buffer containing 14% PEG 8000. The reaction was incubated at 25°C for 2 h and subsequently purified with the Monarch® Spin RNA Cleanup Kit. Reverse transcription was performed on 140 ng of 3’-ligated RNA using Maxima Reverse Transcriptase (Thermo Fisher Scientific) in 1X RT buffer with RNase inhibitors and 1.3 µM RT primer (SB103). The reaction was conducted under the following conditions: 65°C for 5 min, 60°C for 30 min, 65°C for 25 min, and 80°C for 5 min. RNase H and RNase A treatments were performed post-reverse transcription, followed by purification with the Monarch® Spin RNA Cleanup Kit. For 5’ adapter ligation, the cDNA was ligated to a 5’ adapter (SB077) using Thermostable 5’App DNA/RNA Ligase (NEB) in 1X NEB1 buffer with 5 mM MnCl₂. The reaction was incubated overnight at 65°C and purified twice with the Monarch® Spin RNA Cleanup Kit. Library amplification was conducted using indexed primers (SB085 and SB071 or SB072) on 2 ng of cDNA with Q5 High-Fidelity DNA Polymerase (NEB) in 1X Q5 buffer supplemented with 1X GC-rich enhancer. PCR conditions included 25 cycles with an annealing temperature of 60°C. A quarter of the amplified library was resolved on a 10% denaturing PAGE-urea gel. Fragments >85 nt were excised, eluted by soaking overnight (or for 2 h at room temperature), and purified with Monarch® Spin RNA Cleanup Kit columns. Multiplexed libraries were sequenced on an Illumina NextSeq 2000 platform in a 150 bp paired-end configuration.

To map 5’ ends of small tRNA fragments, approximately 100 ng of purified tRNAs from *E. coli* (with or without activated PARIS) or in vitro reactions with purified AriB were end-repaired as described previously [52]. RNA was purified using the RNeasy MinElute Cleanup Kit (QIAGEN) according to the manufacturer’s protocol, with a modified RNA binding step that included 675 µL of 96% ethanol. Purified RNA fragments were eluted in 10 µL of nuclease-free water. Library preparation was performed using the NEBNext® Small RNA Library Prep Set for Illumina® (NEB, #E7330S, USA) following the manufacturer’s instructions. DNA libraries were quantified using a Qubit 2.0 fluorometer (Invitrogen, USA) and assessed for quality using a High Sensitivity DNA chip on an Agilent Bioanalyzer 2100. Libraries were multiplexed and sequenced on an Illumina NextSeq 2000 platform using a 50 bp single-end read mode.

Reads were mapped to a *E. coli* tRNAs and rRNAs list excluding any redundant sequences. Alignment algorithm used was Bowtie2 with default parameters and only uniquely mapped reads were kept for the scatter plots.

### AriB purification

To produce activated AriB, PARIS (AriA-AriB-Strep) from pFR85 and non-tagged Ocr from pBAD were expressed in separate cultures as previously described [12]. Cell cultures were centrifugated and pellets resuspended in buffer A (100mM Tris pH 8.0, 150mM NaCl, 1mM EDTA) containing protease inhibitors cocktail (Abcam, ab271306). Cells were lysed by sonication on ice and cell debris was removed by centrifugation (15.000g, 20 min, 4°C). Cleared filtered lysates of PARIS-and Ocr-expressing cells were mixed and incubated one hour at RT before loading on a 5 mL StrepTrap XT affinity column (Cytiva) equilibrated in buffer A. Activated AriB-Strep was eluted with buffer BXT (IBA Lifesciences). Further purification was done with size exclusion chromatography performed on a Superdex 200 Increase 10/300 column (Cytiva) equilibrated with buffer B (20mM Tris pH 8.0, 250mM NaCl, 1mM DTT). Fractions of interest were combined and concentrated (Amicon Ultra Centrifugal Filter, 30 kDa MWCO).

### DNA template production for toeprint

First, we designed a DNA matrix for T7 *in vitro* transcription that will encode ORF, utilizing all known *E. coli* tRNA species (RST3_all_tRNAs, Table S1). To generate this dsDNA matrix we used two partially overlapping oligos encoding ORF and carrying T7 polymerase binding site. Oligos were annealed with gradual temperature decrease from 98°C to 25°C for 2°C every 2min. To synthesize the missing second strand, DNA Polymerase I, Large (Klenow) Fragment (NEB) was used according to manufacturer’s instructions. After the reaction, DNA templates were purified from excess of dNTPs and polymerase with Monarch PCR & DNA Cleanup Kit (NEB).

### *In vitro* translation and toeprinting analysis

The generated DNA templates were amplified using Q5® High-Fidelity 2X Master Mix (NEB) with T7F (TAATACgACTCACTATAgg) and NV1 (GGTTATAATGAATTTTGCTTATTAAC) primers.

The templates were validated for *in vitro* transcription using MEGAscript® Kit (Thermofisher Scientific) according to manufacturer, supplemented with FAM-11-UTP (0.25 mM). To assess RNA integrity of T7 transcript(s), RNA samples were run on a 6% denaturing polyacrylamide-urea gel electrophoresis, followed by scanning using Typhoon™ FLA 9500. Toeprinting was performed using the PURExpress In Vitro Protein Synthesis Kit (NEB #E6800)and fluorescently labelled probes [53]. The mixture containing 2 µL of solution A, 1 µL of solution B, 0.5 µL of 20 µM FAM-labelled NV1 primer (5ʹ-/FAM/-GGTTATAATGAATTTTGCTTATTAAC-3ʹ, bearing a fluorescein at the 5ʹ-end), 0.2 µL of 40 U/µL Ribolock (Thermofisher Scientific) was treated with either 0.5 µL of 0.6 µg/mL AriB or storage buffer (TAKM_7_ buffer: 50 mM Tris-HCl pH 7.5, 70 mM NH4Cl, 30 mM KCl, and 7 mM MgCl2) for 20 min at 37°C. Thiostrepton (Ths) (50 µM) was used as a control to denote the inhibitory signal at initiation stage. The reactions were started when a DNA template (10 ng) of interest was added to the mixture followed by an additional incubation for 30 min at 37°C. A 1 µL master mix containing 0.1 µL AMV reverse transcriptase (Roche), 0.2 µL 5X AMV buffer (Roche), 0.5 µL of 4 mM dNTP mix (Thermofisher Scientific), 0.2 µL of ultra-pure water was added to the reaction(s) for cDNA synthesis. The AMV reverse transcription reaction was conducted for 15 min at 37°C. The reaction(s) were then terminated using 1 µL of 10 M NaOH, with subsequent incubation for 15 min at 37°C followed by complete neutralization by adding 1 µL of 12 N HCl. The cDNA product was purified using the QIAquick PCR Purification Kit (Qiagen). Fluorescently labelled cDNA products were analysed by capillary gel electrophoresis using a Nanophore-05 Genetic Analyzer (Syntol LLC, Russia). The collected data was processed and visualized using GeneMarker® software (SoftGenetics). The abundance of cDNA fragments scored was denoted by area under the graph that reflects the amount of stalled ribosomes. The ribosome position was deduced by using classical toeprinting principle. The 3’ end of synthesized cDNA product is separated by ∼13-14 nt from the first base of the A-site when reverse transcriptase encounters stalled ribosomes. All experiments were conducted reproducibly at least three times.

### E. coli mutants deficient for tRNA modifications

Strains harboring Δ*mnmA*, Δ*truA*, Δ*truB* or Δ*tcdA* were obtained from the Keio collection. Each deletion was transferred individually to strain MG1655 (FBE051) by P1 transduction and the kanamycin resistance selection marker was then removed using the pE-FLP (AmpR) plasmid able to recombine FRT sites flanking the backbone. pE-FLP was then cured through serial restreaks on LB plates, leading to strains FBE939 (Δ*mnmA*) [37], FD6 (Δ*truA*), FD7 (Δ*truB*) and FD8 (Δ*tcdA*). The integration and then each deletion on the chromosome were checked by PCR with primers followed by Sanger sequencing.

Each strain harbouring the various deletions was transformed, by heat shock at 42°C following standard protocols, with pFR85 (Ptet *ariA-ariB*) or control plasmid pFR66 (Ptet sfGFP) and pFD250 (PPhlF *ocr*) carrying *ocr* under the control of an inducible DAPG PhlF promoter. Then *E. coli* cells carrying these plasmids were grown in LB broth with kanamycin (50 μg/ml) for pFR85 and pFR66 and chloramphenicol (20 μg/ml) for pFD250 (PPhlF *ocr*) overnight at 37°C followed by serial dilutions and spotted on LB agar containing kanamycin and chloramphenicol with induction of Ocr or not by DAPG 50 µM. Strains, primers and plasmids are listed in the Table S1.

### *In vitro* transcripts production and purification

Oligonucleotides containing various tRNA sequences preceded by a 5’extended T7 promoter sequence (CGATTGAGGCCGGTAATACGACTCACTATA) were ordered at IDT or Merk. (oligos Table S1). A 20-cycle PCR was run on 3ng of template oligo with a forward primer corresponding to the 5’extended T7 promoter sequence and a template specific reverse primer antisense of 30nt of the 3’ end of the template oligo in a NEB Q5 HiFi reaction mix. Products were purified with a Macherey-Nagel PCR cleanup column, eluted in 30µl.

0.8-1µg of PCR product was then used as a template for IVT with a HiScribe® T7 High Yield RNA Synthesis Kit. 5 MM of DTT, 10mM NTP (each) and 2µl of T7 RNA Polymerase Mix in a 20µl reaction was carried out overnight at 37°C. IVT product were either PAGE-purified or purified using NEB Monarch RNA cleanup column, following DNase I treatment, and eluted in 20µl RNase-free water.

### *In vitro* AriB nuclease activity

In vitro activity on total enriched small RNA has been done as described in [12]. The activity on *in vitro* transcribed tRNA has been done in 20µl reaction buffer (20mM Tris pH 8.0, 200mM NaCl) containing 100nM (2pmol) purified tRNA and 25nM pure AriB dimer. Magnesium (MgSO_4_) has been added to a final concentration of 1mM except for the experiment of metal-dependency where the final concentration was 0.5mM final alongside other metals (MnCl_2_, CaCl_2_, CoCl_2_, CuSO_4_ and NiCl_2_). The mixture was incubated at 37°C for 20min, otherwise specified, and the reaction stopped with 1:1 v:v of 100% formamide. The samples were incubated 5 min at 92°C before loading 20µl (1pmol) on a preheated 12% denaturing PAGE containing 7M urea. After running the gel at 25W constant for about an hour, the gel was stained with SYBR^TM^ Gold (Invitrogen S11494) for 10min before visualisation on a Bio-Rad Chemidoc MP Imaging System.

### Poly(A) tailing of AriB cleavage products

In a 10µl 1X poly(A) reaction buffer, 2pmol (200nM final) of purified (NEB Monarch RNA cleanup column) AriB cleaved tRNA^Lys(UUU)^ were incubated with 1mM of ATP and 2.5 units of E. coli Poly(A) Polymerase (NEB M0276) for 30min at 37°C. The reaction was stopped with 1:1 v:v of 100% formamide, incubated 5 min at 92°C before loading 1pmol on a preheated 12% denaturing PAGE containing alongside 1pmol of uncut and cut controls.

### Phage infection: EOP assay

The activity of PARIS or PARIS mutant pFD340 (Ptet *ariA ariB* K60A/K64A) was measured by performing efficiency of plaquing (EOP) assays with different phages. *E. coli* BW25113 carrying the control plasmid pFR66 (sfGFP) or the WT PARIS system (pFR85) with pBAD vector encoding tRNAs was grown overnight in LB with kanamycin (50 μg/ml) and ampicillin (100 μg/ml). *E. coli* K-12 MG1655 carrying the double mutant pFD340 (Ptet *ariA ariB* K60A/K64A), the control plasmid pFR66 (sfGFP) or pFR85 (Ptet *ariA ariB* or WT PARIS system) was grown overnight in LB supplemented with 50 μg/ml kanamycin. Bacterial lawns were prepared by mixing 100 μl of a stationary culture with 5 ml of LB + 0.5% agar, and the mixture was poured onto a Petri dish containing LB with kanamycin (50 μg/ml) for pFR85 (Ptet *ariA ariB* or WT PARIS system), pFR66 (sfGFP) and pFD340 (Ptet *ariA ariB* K60A/K64A) and ampicillin (100 μg/ml) in the presence of pBADvector derivatives and supplemented with 1 mM CaCl2. Induction of tRNA genes was achieved by addition in the top agar of the indicated amount of L-Ara. Tenfold serial dilutions of high-titer stock of phages were spotted on each plate and incubated overnight at 37 °C or 5h for T7 phage. Plaque assays were performed in at least two independent replicates.

### Mutant (K60A and K64A) in the cleavage site

The double mutation K60A and K64A in *ariB* was synthesized with primers F747 and F748 (Table S1) and introduced on pFR85 plasmid carrying *ariA* and *ariB* genes with the native promoter and an inducible Ptet promoter upstream through two PCRs with primers F747/LC327 and TG99/F748 (Table S1) of the whole plasmid, followed by Gibson assembly [54]. The resulting plasmid pFD340 (*ariA ariB* K60A/K64A) was introduced into competent *E. coli* K-12 MG1655 cells by heat shock at 42°C, following standard protocols, and plated on LB agar supplemented with 50 μg/ml kanamycin. The construction was verified using Sanger sequencing. Activity of this mutant has been tested by performing EOP assay as described above.

### Structure AF3 and alignment with M5 nuclease

Two sequences of AriB monomer and the sequence of *E. coli* tRNA^Lys(UUU)^ has been loaded on the AF3 server. The predicted template modelling (pTM) score of 0.92 means the overall predicted fold for the complex might be similar to the true structure [55]; and the interface predicted template modelling (ipTM) score de 0.82 suggest a confident high prediction [56]. The ‘align’ command in PyMOL was used to superpose the M5 nuclease (PDB 6z2b) onto AriB. Specifically, chain B of the M5 nuclease was aligned to a selection comprising residues 19-43, 48-51, 80-87, 109-111 and 113-115 of AriB. This led to an rmsd of 0.47 Å over 87 pairs of atoms.

### MS analysis of tRNA^Lys(UUU)^ fragments

5 ug of *in vitro* transcribed *E. coli* tRNA^Lys(UUU)^ was mixed with 1 ul AriB or AriB E26A (16mg/ml) in the reaction buffer (25 mM Tris-HCl, 35 mM NH_4_Cl, 15 mM KCl, 5 mM MgCl_2_, 0.5 mM DTT, pH = 7.5). The reaction mixture was incubated at 37°C for 30 min. Purification of tRNA and fragments from reaction was performed with «Lira» (Biolabmix, Russia, LR-100). After, re-extracted tRNA was mixed with 20U RNase T1 (Thermo Fisher Scientific, USA) in 30 mM NH_4_Ac and incubated at 37°C for 30 min. Reaction aliquote was mixed with 2,5-dihydroxybenzoic acid solution (40 mg/mL in 30% acetonitrile, 0.5% TFA) and analyzed by mass-spectrometry using UltrafleXetreme MALDI-TOF*/*TOF (Bruker Daltonik, Germany) equipped with Nd laser. The average MH+ molecular ions of RNase T1-treated tRNA were measured in linear mode; the accuracy of average mass peak measurement was within 1 Da

