## SupplementaryData for "Specificity and Mechanism of tRNA cleavage by the AriB Toprim nuclease of the PARIS bacterial immune system": SupplentaryData.pdf

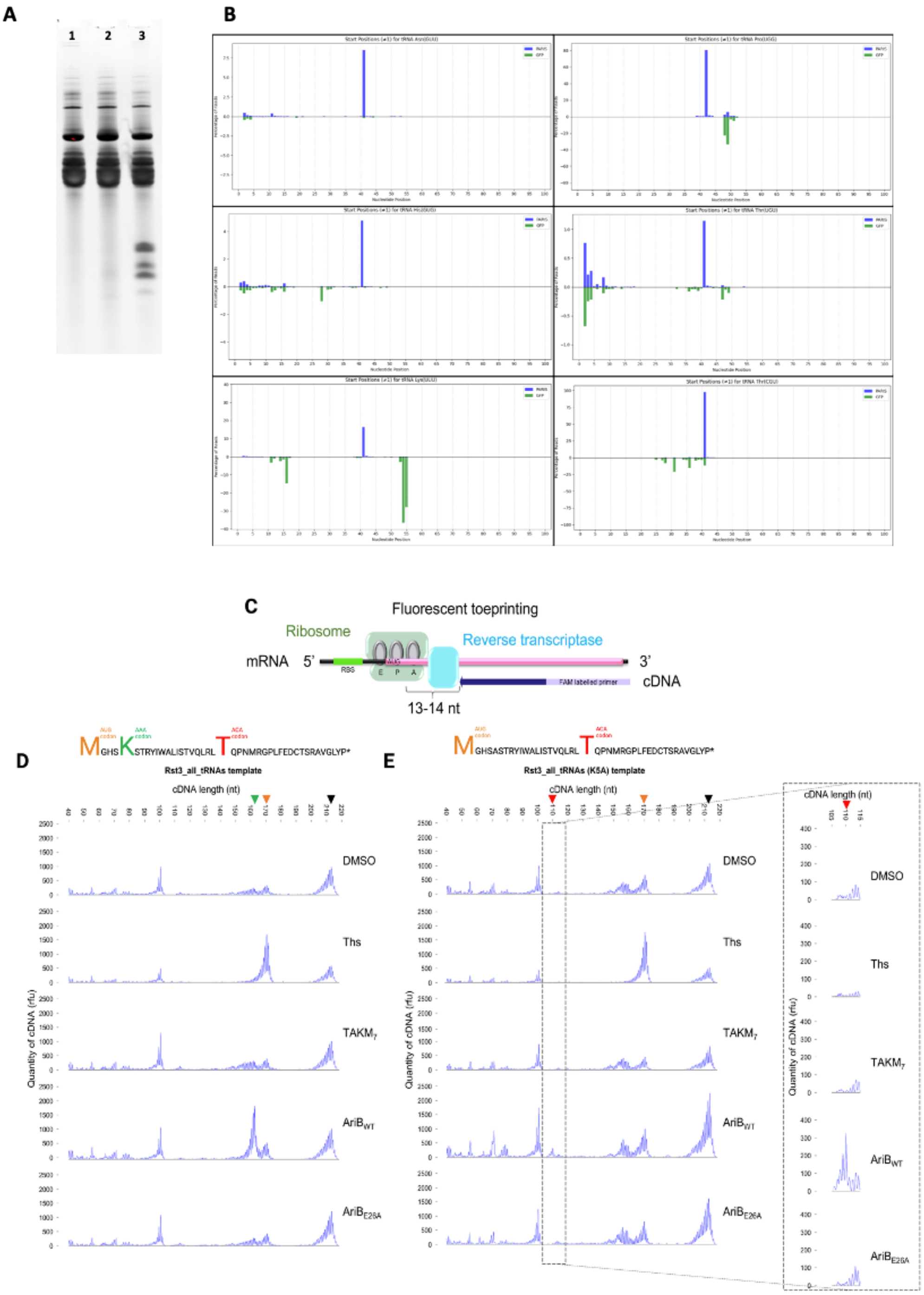


**Supplementary figure S1. (extended figure 1)**

**A.** Visualization on denaturing PAGE of the samples that have been used for sequencing; 1, Ocr/GFP; 2, Ocr/PARIS; 3, AriB cleavage *in vitro*. **B.** Histogram for detection of cleavage site by 5' end counting of sequenced reads. The histograms display the normalized abundance of the 5' of reads for the specified tRNA in both samples GFP control and PARIS. The x-axis represents the position along the tRNA sequence, while the y-axis shows the relative abundance of the 5'sites in the reads for that tRNA. **C.** A general principle of fluorescent toeprinting assay. **D-E** Distribution of cDNA fragment lengths after *in vitro* translation in the presence of AriB WT or E26A Toprim mutant, $TAKM_7$ is a control reaction with buffer. The black arrow indicates full length mRNA *rst3m_all_tRNAs*, green arrow highlights the position of the lysine AAA codon, while red arrow indicates the position of the threonine ACA codon. Positions of the ribosome are shown relative to the A-site. In the presence of AriB WT ribosomes are stalled on lysine (AAA) or threonine (ACA) codons of *rst3m_all_tRNAs* and *rst3m_all_tRNAs K5A* mRNA templates respectively.

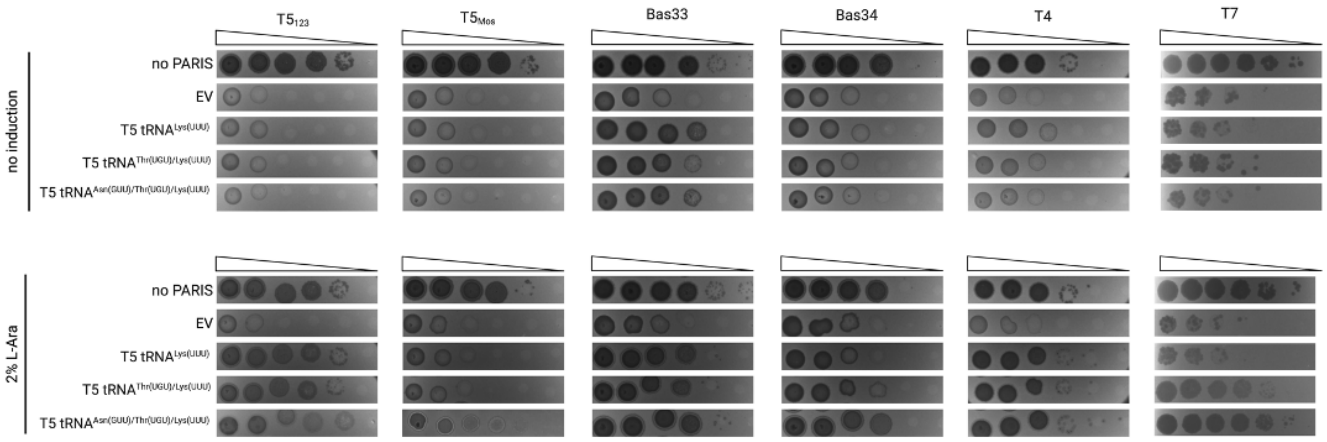


**Supplementary figure S2. (extended figure 2)**
EOP assay showing PARIS culture infection efficiency for the phages $T5_{123}$, $T5_{Mos}$, Bas33, Bas34, T4, T7 in conditions of expression of the indicated sets of T5 tRNAs. tRNAs were expressed from pBAD in conditions of 2% L-arabinose induction.

**A**

| | 8 | 16 | 17 | 20 | 32 | 34 | 37 | 39 | 46 | 47 | 54 | 55 |
|---|---|---|---|---|---|---|---|---|---|---|---|---|
| Lys(UUU) | | D | D | D | | mnm$^5$s$^2$U | ct$^6$A | Y | m$^7$G | acp$^3$U | m$^5$U | Y |
| Thr(UGU) | | | | D | Cm | cmo$^5$U | t$^6$A | | m$^7$G | | | Y |
| Asn(GUU) | s$^4$U | D | | D | | Q | ct$^6$A | Y | m$^7$G | | m$^5$U | Y |
| Thr(CGU) | | D | D | D | | | m$^6$t$^6$A | | m$^7$G | | m$^5$U | Y |

- **s$^4$U** 4-thiouridine
- **D** Dihydrouridine
- **Cm** 2′-O-methylcytidine
- **mnm$^5$s$^2$U** 5-methylaminomethyl-2-thiouridine
- **cmo5U** uridine 5-oxyacetic acid
- **Q** Queuosine
- **t$^6$A** N6-threonylcarbamoyladenosine
- **ct$^6$A** Cyclic N6-threonylcarbamoyladenosine
- **m$^6$t$^6$** N6-methyl-N6-threonylcarbamoyladenosine
- **Y** Pseudouridine
- **m$^7$G** 7-methylguanosine
- **acp$^3$U** 3-(3-amino-3-carboxypropyl)uridine
- **m$^5$U** 5-methyluridine

**B**

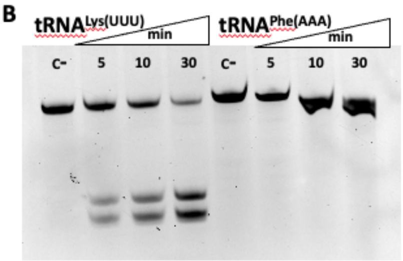


**C**

```
                     10        20        30  anticodon   40-41   50
Lys(UUU) GGGUCGUUA--GCUCAGUUGGUAGAGCAGUUGACUUUUAAUCAAUUGGUCGCAGGUUCGAAUCCUGCACGACCCACCA
Thr(UGU) --GCCGACUUAGCUCAGUAGGUAGAGCAACUGACUUGUAAUCAGUAGGUCACCAGUUCGAUUCCGGUAGUCGGCACCA
Asn(GUU) -GUCCUCUGUAGUUCAGUCGGUAGAACGGGTGACUGUUAAUCCGUAUGUCACUGGUUCGAGUCCAGUCAGAGGAGCCA
Thr(CGU) --GCCGAUAUAGCUCAGUUGGUAGAGCAGCGCAUUCGUAAUGCGAAGGUCGUAGGUUCGACUCCUAUUAUCGGCACCA
             *      * ***** ****** *     * *  ****      ***    ****** ***          ***

                                          anticodon
Ec Lys(UUU) GGGUCGUUAGCUCAGUUGGU--AGAGCAGUUGACUUUUAAUCAAUUGGUCGCAGGUUCGAAUCCUGCACGACCCACCA
T5 Lys(UUU) GGGUUGCUAGCUCAACUGGUUUAGAGCACUGGUCUUUUAAACCAUAGGUUACAGGUUCGAGUCCUGUGCAACCCACCA
            **** * *******  ****  ****** * * ******* * ** ***  ********* *****  * ********
Ec Thr(UGU)   GCCGACUUAGCUCAGUAGGUAGAGCAACUGACUUGUAAUCAGUAGGUCACCAGUUCGAUUCCGGUAGUCGGCACCA
T5 Thr(UGU)   GCUCCUAAAGCAUUGUUGGUGAUGCAGUUGCCUUGUAAGCAUCUGAAC-CGGGUUCGAUUCCUAGUGGGAGCACCA
              **      ***   ** ***   ***  ** ******* **   *  * *  **********    *   ******
Ec Asn(GUU) UCCUCUGUAGUUC-AGUCGGU-AGAACGGCGGACUGUUAAUCCGUAUGUCACUGGUUCGAGUCCAGUCAGAGGAGCCA
T5 Asn(GUU) GGUUCCUUAGCUCUAAUUGGUUAGAGCGGCAUCUUGUUAAGUUGAGGGUUGCUGGUUCGAAUCCAGCAGGAACCGCCA
               **  *** ** * * *** *** ****    ******   *   ** ********* *****   **    ****
```

**D**

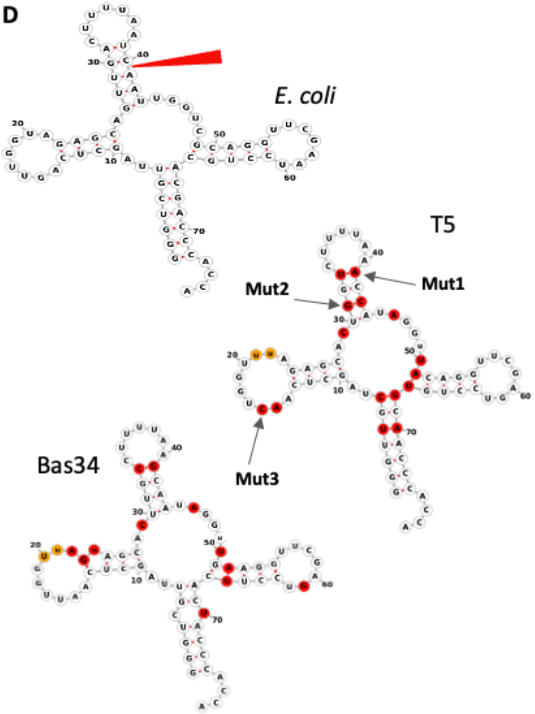


**Supplementary figure S3. (extended figure 3.)**

**A.** Visualization of modifications on selected *E. coli* tRNAs corresponding to Lys(UUU), Thr(UGU), Asn(GUU) and Thr(CGU) using the Modomics server [1]. **B.** Time course of AriB activity on *in vitro* transcripts corresponding to *E. coli* tRNA$^{Lys(UUU)}$ and tRNA$^{Phe(AAA)}$. **C. Top.** Multiple Sequence alignment by CLUSTALW of tRNA$^{Lys(UUU)}$, tRNA$^{Thr(UGU)}$, tRNA$^{Asn(GUU)}$ and tRNA$^{Thr(CGU)}$. The cleavage site is indicated by a red mark; the common sequence $U_{36}AAUCA_{41}$ found in tRNA$^{Lys(UUU)}$ and tRNA$^{Thr(UGU)}$ is framed in blue. **Bottom.** Sequence alignment between *E. coli* and T5 corresponding tRNA. **D.** Comparison of E. coli, T5, and Bas34 tRNA$^{Lys(UUU)}$. Substitutions are indicated in red and insertions in yellow.

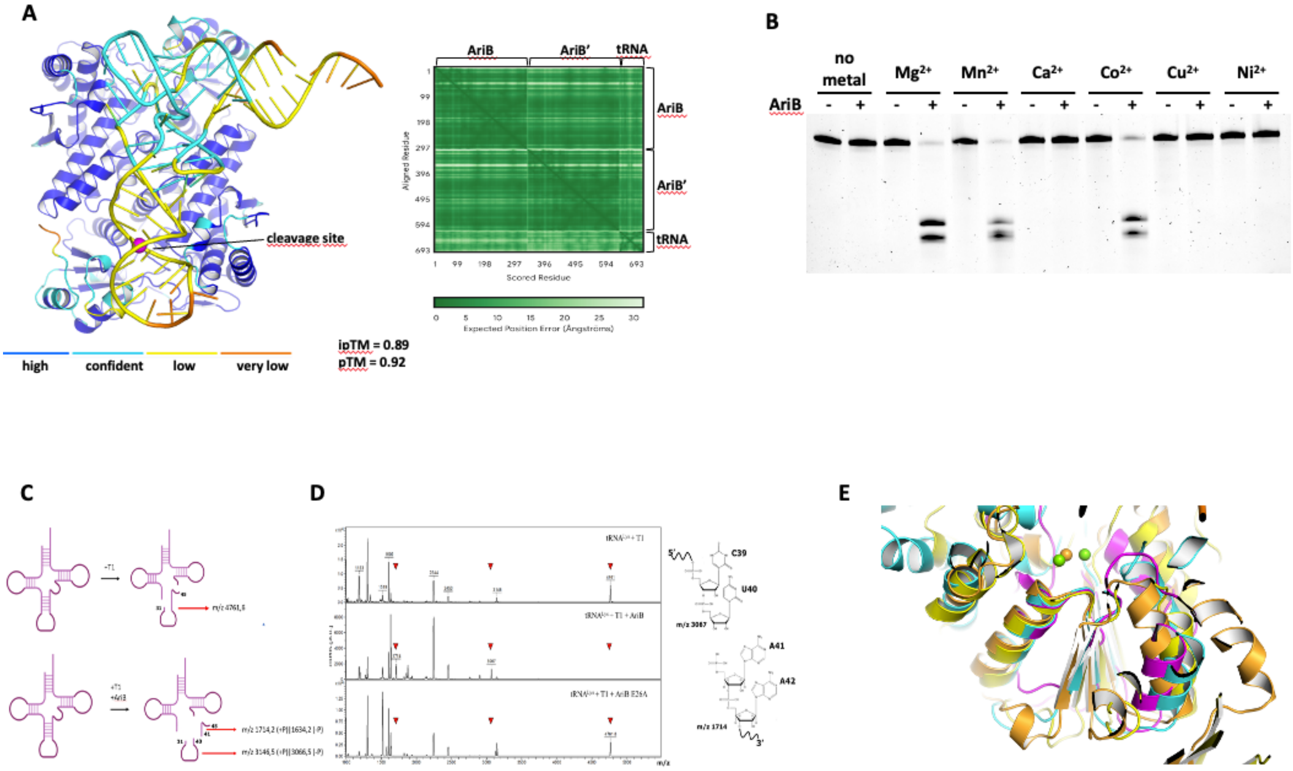


**Supplementary figure S4. (extended figure 4.) AriB structure, metal dependency and cleavage products.**
**A.** AF3 structure with high confidence as measured by predicted Local Distance Difference Test (pLDDT) **B.** Metal dependency of *in vitro* AriB activity on *E. coli* tRNA$^{Lys(UUU)}$ transcript. 100nM of RNA is incubated with 0.5mM of metal ions in presence or not of 25nM of AriB. **C.** T1 RNAse cleavage of the intact *E. coli* tRNA$^{Lys(UUU)}$ produces a 14-bp fragment with *m/z*=4761,6; if the *E. coli* tRNA$^{Lys(UUU)}$ is nicked by AriB, T1 RNAse treatment produces two sub-fragments. Depending on the -$PO_4$ group retention at the 3' or 5' end of the cleavage products, these sub-fragments will have distinct *m/z* values. **D.** MS analysis of tRNA$^{Lys(UUU)}$ T1 RNAse cleavage products. The $MH^+$ ions at *m/z*=1714, *m/z*=3067, corresponding to the sub-fragments of the original *m/z*=4761,6 fragment, could be detected only in reactions with AriB WT, but not AriB E26A, and support formation of the 3'-OH and 5'-$PO_4$ AriB cleavage products (on the right). **E.** Superposition of AriB (cyan), M5 (magenta), GajA (yellow) and TsOLD (orange) nuclease active sites. The magenta and orange spheres represent the metal ions of the corresponding-coloured structures.

**Supplementary material (PSE file). Structure of AriB bound to *E. coli* tRNA$^{Lys(UUU)}$ predicted by AF3.**

AriB monomers are respectively in yellow gold (with the active catalytic site) and blue (with the unactive catalytic site). The conserved Toprim residues are labelled with red sticks and the two lysine residues K60/K64 with pale blue sticks. The RNA is pale green with the cleavage site ($C_{40}A_{41}$) in purple sticks. In orange sticks are the nucleotides facing the unactive catalytic site. Magnesium ions are represented as blue spheres; in each catalytic site one is predicted whereas the two others are predicted interacting with the tRNA.

**Supplementary Table S1:**

<table>
<tr><th>E. coli strain</th><th colspan="2">Comment</th><th>Source</th></tr>
<tr><td>BW25113</td><td colspan="2">E. coli K12 F– Δ(araD-araB)567 ΔlacZ4787(::rrnB-3) λ– rph-1 Δ(rhaD-rhaB)568 hsdR514</td><td>Lab stock</td></tr>
<tr><td>XL1-Blue</td><td colspan="2">E. coli K12 recA1 endA1 gyrA96 thi-1 hsdR17 supE44 relA1 lac [F'proAB lacIqZΔM15 Tn10], TetR</td><td>Evrogen</td></tr>
<tr><td>MG1655</td><td colspan="2">E. coli K12 F-, λ-, ilvG-, rfb-50, rph-1</td><td>Lab stock</td></tr>
<tr><td>MG1655 (FBE051)</td><td colspan="2">E. coli K12 F-, λ-, ilvG-, rfb-50, rph-1</td><td>Lab stock</td></tr>
<tr><td>FBE939</td><td colspan="2">MG1655 (FBE051)::ΔmnmA</td><td>[2]</td></tr>
<tr><td>FD6</td><td colspan="2">MG1655 (FBE051)::ΔtruA</td><td>This study</td></tr>
<tr><td>FD7</td><td colspan="2">MG1655 (FBE051)::ΔtruB</td><td>This study</td></tr>
<tr><td>FD8</td><td colspan="2">MG1655 (FBE051)::ΔtcdA</td><td>This study</td></tr>
<tr><td>Phage</td><td colspan="2">Comment</td><td>Source</td></tr>
<tr><td>T7</td><td colspan="2"></td><td>Lab stock</td></tr>
<tr><td>T5</td><td colspan="2"></td><td>Lab stock</td></tr>
<tr><td>T4</td><td colspan="2"></td><td>Lab stock</td></tr>
<tr><td>$T5_{Mos}$</td><td colspan="2">T5 with 30661-38625 deletion</td><td rowspan="2">[3]</td></tr>
<tr><td>$T5_{123}$</td><td colspan="2">T5 with 29191-32442 deletion</td></tr>
<tr><td>Bas33<br>HildyBeyeler</td><td colspan="2"></td><td>[4]</td></tr>
<tr><td>Bas34<br>SelmaRatti</td><td colspan="2"></td><td>[4]</td></tr>
<tr><td>Plasmid Name</td><td>Plasmid Map</td><td>Comment</td><td>Source</td></tr>
<tr><td>pE-FLP</td><td>https://benchling.com/s/seq-ZSAOyudBu7l6aE1fA7ZA?</td><td>FRT sites recombination</td><td>[5]</td></tr>
</table>

| | | | |
|---|---|---|---|
| | m=slm-iBlwknIMfOyMDlrkDObX | | |
| pFR66 | https://benchling.com/s/seq-UjYoHgnFK8K3YMeLF3LO?m=slm-iVxvAYV8bipLZ5NYOQp0 | sfGFP, Tet promoter, $Kan^R$ | [6] |
| pFR85 (PARIS) | https://benchling.com/s/seq-P0S4QJUYX0D54AVNp81e?m=slm-DWv2SMSVbNQMxjbonDYa | pFR66 with PARIS-2 from *E.coli* B185, native PARIS-2 promoter + Ptet promoter, $Kan^R$ | [6] |
| pBAD T5 $tRNA^{Lys}$ = pFD287 | https://benchling.com/s/seq-vMWoZV7cTRTJmxHolzoL?m=slm-lwz3eDeiC18WokES0QGM | T5 $tRNA^{Lys}$ (UUU) cloned together with 20 bp upstream and downstream under control of *araBAD* promoter, $Amp^R$ | [7] |
| pBAD T5 $tRNA^{AsnThrLys}$ | https://benchling.com/s/seq-Zbfh22XXcmpU8KJ0s4X7?m=slm-6dQamFwN7Cp3yG9jb243 | T5 $tRNA^{Lys}$ (UUU) $tRNA^{Asn}$ (GTT) $tRNA^{Thr}$ (TGT) cloned together with 20 bp upstream and downstream under control of *araBAD* promoter, $Amp^R$ | This study |
| pFD250 | https://benchling.com/s/seq-3Y0xK3fj0qRzXKjSqkrH?m=slm-WXcGNkiVOWzJvLAupoOb | pFD200 with T7 *0.3* gene encoding Ocr under the control of $P_{PhlF}$ promoter, $Cm^R$ | [7] |
| pBAD Ocr | https://benchling.com/s/seq-Mcb9hM3HgRdwZo0A3YP | Phage T7 Ocr cloned under control of *araBAD* promoter, $Amp^R$ | [8] |

| | | | |
|---|---|---|---|
| | 2?m=slm-2UNmUVUYQHtIqvg3LuOL | | |
| pFD340 | https://benchling.com/s/seq-9C912jwctAX5RxHbdmGq?m=slm-m4370F7EVbTWstKu8m9a | pFR85 with AriB K60A K64A double mutation, $Kan^R$ | This study |
| RST3_all_tRNAs | https://benchling.com/s/seq-uOpcdgLuXeP3rkgb5JUr?m=slm-RqbFbqhZnoXx0u5N1sdh | DNA matrix for in vitro transcription mRNA for toe print | This study |
| *RST3_all_tRNAs_noK* | https://benchling.com/s/seq-UsCCEcRw6V5ugConUjPW?m=slm-1E143rUU7HwQA0Jhl3OQ | DNA matrix for in vitro transcription mRNA without LysU codon for toe print | This study |

| **Primer Name** | **Sequence (5'->3')** | **Template** | **Comment** |
|---|---|---|---|
| pBAD_For | ATGCCATAGCATTTTTATCC | pBAD18 | pBAD sequencing primers |
| pBAD_Rev | GATTTAATCTGTATCAGGCTG | | |
| pBAD256b_F | TTTTTATAACCTCCTTAGAGCTCG | pBAD T5 $tRNA^{Lys}$ | Amplification of pBAD with T5 $tRNA^{Lys}$ backbone for Gibson Assembly |
| 256b_tRNA_R | AAGTATATAATGGTTACTAAGGGTTG | | |
| 256_T5_Lys+Asn_F | TTTTTGGGCTAGCGAATTCGAGCTCACTCTAGTTCTGACTTATATGG | T5 genomic DNA | Amplification of T5 $tRNA^{Asn}$ for Gibson Assembly |
| 256_T5_Lys+AsnThr_R | GATGATCCCGTTATTTCAACACTTAACTCTACACAGAATTTG | | |
| 256_T5_Thr_F | GTTGAAATAACGGGATC | | Amplification of T5 $tRNA^{Thr}$ for Gibson Assembly |
| 256_T5_Lys+ThrAsn_R | CAACCCTTAGTAACCATTATATACTTGCAAATGAATTTTTAAG | | |
| Toe_print_alltRNA_F | ACTAATACGACTCACTATAGGGCTTAAGTATAAGGAGGAAAACATATGGGACACTCAAAATCGACGAGGTACATATGGGCATTGATCAGCACTGTACAACTGCGTCTAACACAGCCGAACATGCGG | | For toe print DNA matrix synthesis |
| Toe_print_alltRNA_R | GGTTATAATGAATTTTGCTTATTAACGATAGAATTCTATCACATTCTTGATTCTTATTAGGGATATAACCCGACGGCTCTGGAGGTGCAGTCTTCGAAGAGTGGGCCCCGCATGTTCGGCTGTGTTAGACGC | | |
| Toe_print_K5A_tRNA_F | ACTAATACGACTCACTATAGGGCTTAAGTATAAGGAGGAAAACATATGGGACACTCAGCATCGACGAGGTACATATGGGCATTGATCAGCACTGTACAACTGCGTCTAACACAGCCGAACATGCGG | | |

| | | | |
|---|---|---|---|
| Toe_print_noLysQ_noThrU_F | ACTAATACGACTCACTATAGGGCTTAAGTATAAGGAGGAAAACATATGGGACACTCAGCATCGACGAGGTACATATGGGCATTGATCAGCACTGTACAACTGCGTCTACAGCCGAACATGCGG | | |
| Toe_print_noLysQ_noThrU_R | CTGCGTCTACAGCCGAACATGCGGGGCCCACTCTTCGAAGACTGCACCTCCAGAGCCGTCGGGTTATATCCCTAATAAGAATCAAGAATGTGATAGAATTCTATCGTTAATAAGCAAAATTCATTATAACC | | |
| T7F | TAATACGACTCACTATAGG | | For toe print mRNA matrix synthesis |
| NV1 | GGTTATAATGAATTTTGCTTATTAAC | | For toe print cDNA synthesis |
| Ecoli_Lys_transcript_F | ATTAATACGACTCACTATGGGTCGTTAGCTCAGTTGGTAGAGCAGTTGACTTTTAATCAATTGGTCGCAGGTTCGAATCCTGCACGACCCACCA | | For in vitro tRNA transcript synthesis used in the MS approach |
| Ecoli_Lys_ transcript _R | TGGTGGGTCGTGCAGGATTCGAACCTGCGACCAATTGATTAAAAGTCAACTGCTCTACCAACTGAGCTAACGACCCATAGTGAGTCGTATTAAT | | |
| F747_F | AGCATCTGACGCGGCTATGTCTGCGAATAATAGGTTAAAAATAGAAACCATTCATTCG | pFR85 | Double mutation in ariB-K60A K64A of pFR85 (PARIS) using Gibson assembly leading to pFD340 |
| LC327_R | CGCCTTCTTGACGAGTTCTT | | |
| TG99_F | GGATCTATCAACAGGAGTC | pFR85 | |
| F748_R | CCTATTATTCGCAGACATAGCCGCGTCAGATGCTTTTATATCTTGTGC | | |
| F763_F | GATAGCGAAGGTAGCGTGC | truA | Checking the integration and then the deletion of *truA* |
| F764_R | CGAATTGGGATTACTGAACAAC | | |
| F765_F | GCATGATGACCACCGTTTCC | truB | Checking the integration and then the deletion of *truB* |
| F766_R | CTGCAGGTGGTTGATCTGTG | | |
| F767_F | GTCAGTACGAACTGCGTCTG | tcdA | Checking the integration and then the deletion of *tcdA* |
| F768_R | CAAAGAGTGATGTGGATGCG | | |
| CR01_Lys_PCRtempl | CGATTGAGGCCGGTAATACGACTCACTATAGGGTCGTTAGCTCAGTTGGTAGAGCAGTTGACTTTTAATCAATTGGTCGCAGGTTCGAATCCTGCACGACCCACCA | | Oligo as template to generate the IVT template |
| CR01_Lys_PCRtempl | CGATTGAGGCCGGTAATACGACTCACTATAGGGTCGTTAGCTCAGTTGGTAGAGCAGTTGACTTTTAATCAATTGGTCGCAGGTTCGAATCCTGCACGACCCACCA | | Oligo as template to generate the IVT template |
| CR03_Phe_PCRtempl | CGATTGAGGCCGGTAATACGACTCACTATAGCCCGGATAGCTCAGTCGGTAGAGCAGGGGATTGAAAATCCCCGTGTCCTTGGTTCGATTCCGAGTCCGGGCACCA | | Oligo as template to generate the IVT template |
| CR05_T5Lys_PCRtempl | CGATTGAGGCCGGTAATACGACTCACTATAGGGTTGCTAGCTCAACTGGTTTAGAGCACTGGTCTTTTAAACCATAGGTTACAGGTTCGAGTCCTGTGCAACCCACCA | | Oligo as template to generate the IVT template |
| CR07_Lys-M1_PCRtempl | CGATTGAGGCCGGTAATACGACTCACTATAGGGTCGTTAGCTCAGTTGGTAGAGCAGTTGTCTTTTAAACAATTGGTCGCAGGTTCGAATCCTGCACGACCCACCA | | Oligo as template to generate the IVT template |
| CR09_Lys-M2_PCRtempl | CGATTGAGGCCGGTAATACGACTCACTATAGGGTCGTTAGCTCAGTTGGTAGAGCAGTGGACTTTTAATCCATTGGTCGCAGGTTCGAATCCTGCACGACCCACCA | | Oligo as template to generate the IVT template |
| CR11_Lys-M3_PCRtempl | CGATTGAGGCCGGTAATACGACTCACTATAGGGTCGTTAGCTCAGCTGGTAGAGCAGTTGACTTTTAATCAATTGGTCGCAGGTTCGAATCCTGCACGACCCACCA | | Oligo as template to generate the IVT template |
| CR13_LysM1M2_PCRtempl | CGATTGAGGCCGGTAATACGACTCACTATAGGGTCGTTAGCTCAGTTGGTAGAGCAGTGGTCTTTTAAACCATTGGTCGCAGGTTCGAATCCTGCACGACCCACCA | | Oligo as template to generate the IVT template |

| | | | |
|---|---|---|---|
| CR15_ThrUGU_PCRtempl | CGATTGAGGCCGGTAATACGACTCACTATAGCCGACTTAGCTCAGTAGGTAGAGCAACTGACTTGTAATCAGTAGGTCACCAGTTCGATTCCGGTAGTCGGCACCA | | Oligo as template to generate the IVT template |
| CR17_Asn_PCRtempl | CGATTGAGGCCGGTAATACGACTCACTATAGTCCTCTGTAGTTCAGTCGGTAGAACGGCGGACTGTTAATCCGTATGTCACTGGTTCGAGTCCAGTCAGAGGAGCCA | | Oligo as template to generate the IVT template |
| CR19_AsnM2_PCRtempl | CGATTGAGGCCGGTAATACGACTCACTATAGTCCTCTGTAGTTCAGTCGGTAGAACGGCTGACTGTTAATCAGTATGTCACTGGTTCGAGTCCAGTCAGAGGAGCCA | | Oligo as template to generate the IVT template |
| CR21_T5Lys-EcCS_PCRtempl | CGATTGAGGCCGGTAATACGACTCACTATAGGGTTGCTAGCTCAACTGGTTTAGAGCACTTGACTTTTAATCAATAGGTTACAGGTTCGAGTCCTGTGCAACCCACCA | | Oligo as template to generate the IVT template |
| CR23_T5Lys-EcAC-DL_PCRtempl | CGATTGAGGCCGGTAATACGACTCACTATAGGGTTGCTAGCTCAGTTGGTAGAGCAGTTGACTTTTAATCAATAGGTTACAGGTTCGAGTCCTGTGCAACCCACCA | | Oligo as template to generate the IVT template |
| CR25_GlnUUG_PCRtempl | CGATTGAGGCCGGTAATACGACTCACTATATGGGGTATCGCCAAGCGGTAAGGCACCGGTTTTTGATACCGGCATTCCCTGGTTCGAATCCAGGTACCCCAGCCA | | Oligo as template to generate the IVT template |
| SB116_Met_PCRtempl | CGATTGAGGCCGGTAATACGACTCACTATAGGCTACGTAGCTCAGTTGGTTAGAGCACATCACTCATAATGATGGGGTCACAGGTTCGAATCCCGTCGTAGCCACCA | | Oligo as template to generate the IVT template |
| SB117_ThrCGU_PCRtempl | CGATTGAGGCCGGTAATACGACTCACTATAGCCGATATAGCTCAGTTGGTAGAGCAGCGCATTCGTAATGCGAAGGTCGTAGGTTCGACTCCTATTATCGGCACCA | | Oligo as template to generate the IVT template |
| SB109as-Phe-3e | TGGTGCCCGGACTCGGAATCGAACCAAGGA | | reverse primer for PCR to generate IVT template |
| SB110as-T5Lys-3e | TGGTGGGTTGCACAGGACTCGAACCTGTAA | | reverse primer for PCR to generate IVT template |
| SB111as-ThrU-3e | TGGTGCCGACTACCGGAATCGAACTGGTGA | | reverse primer for PCR to generate IVT template |
| SB112as-AsnT-3e | TGGCTCCTCTGACTGGACTCGAACCAGTGA | | reverse primer for PCR to generate IVT template |
| SB113as-Gln1-3e | TGGCTGGGGTACCTGGATTCGAACCAGGGA | | reverse primer for PCR to generate IVT template |
| SB114as-MetT-3e | TGGTGGCTACGACGGGATTCGAACCTGTGA | | reverse primer for PCR to generate IVT template |
| SB115as-ThrW-3e | TGGTGCCGATAATAGGAGTCGAACCTACGA | | reverse primer for PCR to generate IVT template |
| T7promIVTsh2 | CGATTGAGGCCGGTAATACGACTCACTATA | | forward primer for PCR to generate IVT template |
| SB103_Read2-3end | GTGTGCTCTTCCGATCT | | RT primer |
| SB104_ligRTlig_3'Ad-Hyb | /5Phos/NNNNAGAUCGGAAGAGCACA/3AmMO/ | | 5’Phosphate to be turn into 5’App with Ligase I |
| SB077_TGIRT3end4aden | /5Phos/NNNNNNNN CTGTCTCTTATACACATCTGAC/3AmMO/ | | 5’Phosphate to be turn into 5’App with Ligase I |
| SB107_Read2-3end_PS-LNA | T*G*T*G*C*TCTTCCGAT+C+T | | * Indicates phosphothiorate bonds. +N are LNW bases |
| SB071_NEBNforIllum-i01 | CAAGCAGAAGACGGCATACGAGAT CGTGAT GTGACTGGAGTTCAGACGTGTGCTCTTCCGATC*T | | * Indicates phosphothiorate bonds |
| SB072_NEBNforIllum-i02 | CAAGCAGAAGACGGCATACGAGAT ACATCG GTGACTGGAGTTCAGACGTGTGCTCTTCCGATC*T | | * Indicates phosphothiorate bonds |
| SB085_P5-i503-NTread1 | AATGATACGGCGACCACCGAGATCTACAC CCTATCCT TCGTCGGCAGCGTCAGATGTGTATAAGAGACA*G | | * Indicates phosphothiorate bonds |
